# Base Editor Scanning Reveals Activating Mutations of DNMT3A

**DOI:** 10.1101/2023.04.12.536656

**Authors:** Emma M. Garcia, Nicholas Z. Lue, Jessica K. Liang, Whitney K. Lieberman, Derek D. Hwang, James Woods, Brian B. Liau

## Abstract

DNA methyltransferase 3A (DNMT3A) is a *de novo* cytosine methyltransferase responsible for establishing proper DNA methylation during mammalian development. Loss-of-function (LOF) mutations to DNMT3A, including the hotspot mutation R882H, frequently occur in developmental growth disorders and hematological diseases, including clonal hematopoiesis (CH) and acute myeloid leukemia (AML). Accordingly, identifying mechanisms that activate DNMT3A is of both fundamental and therapeutic interest. Here, we applied a base editor mutational scanning strategy with an improved DNA methylation reporter to systematically identify DNMT3A activating mutations in cells. By integrating an optimized cellular recruitment strategy with paired isogenic cell lines with or without the LOF hotspot R882H mutation, we identify and validate three distinct hyperactivating mutations within or interacting with the regulatory ADD domain of DNMT3A, nominating these regions as potential functional target sites for pharmacological intervention. Notably, these mutations are still activating in the context of a heterozygous R882H mutation. Altogether, we showcase the utility of base editor scanning for discovering functional regions of target proteins.

**Synopsis:** 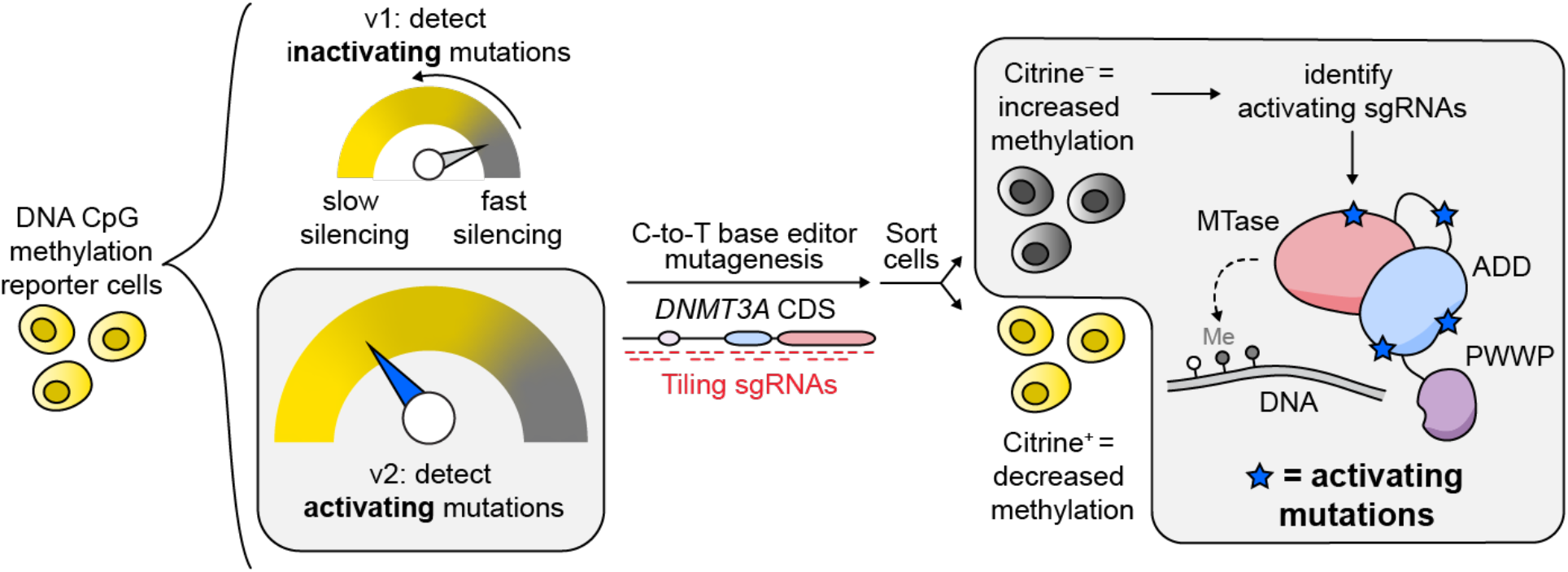

Using base editor mutagenesis and a DNA methylation reporter optimized to find activating mutations, we identify novel hyperactivating mutations in DNMT3A that suggest new mechanisms of allosteric control.

## Introduction

DNMT3A is one of the most highly mutated proteins in hematopoietic disorders, with more than 20% of patients with acute myeloid leukemia (AML) displaying an alteration in *DNMT3A*.^1^ About 65% of these occur at the hotspot R882 codon, which is most frequently mutated to histidine (R882H).^1^ The R882H mutation has been shown to have a dominant-negative effect on DNMT3A activity.^1–4^ Of the non-R882H mutations, a majority have also been shown to be LOF.^5^ DNMT3A LOF mutations also occur in developmental disorders, such as Tatton-Brown–Rahman Syndrome.^6^ Mutations to DNMT3A in hematopoietic disorders are typically heterozygous and associated with a worse prognosis for AML patients.^1,7^ In addition, *DNMT3A* mutations are frequently observed in pre-leukemic hematopoietic stem cell (HSC) populations, in which they promote self-renewal.^8^ Taken together, these observations indicate that *DNMT3A* serves as a tumor suppressor, where mutations drive clonal expansion of HSCs that can eventually lead to AML. Because these disorders all involve LOF mutations in *DNMT3A*, determining mechanisms that may activate DNMT3A or restore its activity could inspire strategies to therapeutically correct disease-associated loss of DNMT3A activity.

DNMT3A has three protein domains: the Pro-Trp-Trp-Pro (PWWP) domain, the ATRX-DNMT3-DNMT3L (ADD) domain, and the catalytic methyltransferase (MTase) domain, along with a disordered N-terminus.^9^ Crystallographic and biochemical data^10,11^ support an allosteric mechanism by which the ADD domain regulates enzyme activity; in the absence of unmodified histone 3 lysine 4 (H3K4me0), DNMT3A adopts an autoinhibited conformation where the ADD domain blocks the binding of the DNA substrate to the catalytic domain. When the ADD domain binds to H3K4me0, it undergoes a conformational change, revealing the DNA binding site to relieve autoinhibition and increase the activity of DNMT3A.^10^ Profiling mutations^5^ that can perturb these regulatory mechanisms, beyond those found clinically, may provide further insight into strategies to activate DNMT3A.

Motivated by these prospects, we conducted *in situ* mutational scanning on *DNMT3A* using base editing.^12^ Base editing is a form of precision genome editing that has enabled the direct mutational scanning of a variety of endogenous proteins.^13–16^ Base editing has the advantage of targeting the native gene, avoiding the confounding effects of overexpressing mutants of a protein of interest. By coupling a fluorescence-based DNA methylation reporter system with base editing to generate a library of mutants, we previously identified a wide variety of LOF mutations in *DNMT3A*, and highlighted the importance of the PWWP domain for enzyme activity, particularly in its ability to bind to DNA.^12^ However, this study only identified a single activating mutation to DNMT3A, which disrupts the known autoinhibitory ADD–MTase interaction.^10^ Whether additional activating mechanisms exist remains an open question.

To explore potential novel activating mutations, in this study we employ base editor scanning coupled to an optimized DNA methylation reporter assay specifically tailored to discover activating *DNMT3A* mutations. By performing the screen in isogenic cell lines with or without the hotspot LOF DNMT3A^R882H^ mutation, we identify mutations that both boost the activity of DNMT3A^WT^ and restore activity in the context of DNMT3A^R882H^. By further validating these mutations biochemically, we nominate the ADD domain and the ADD–MTase interface of DNMT3A as promising functional sites for future pharmacologic modulation to potentiate DNMT3A activity.

## Results

### Development of a screening approach for discovery of activating mutations

To identify activating DNMT3A mutations, we sought to adapt our previously reported base editor scanning approach coupled to a fluorescence-based reporter of DNA methylation.^12,17^ In this system, we employed a reporter construct that contains TetO sites upstream of a promoter controlling the expression of citrine (Figure 1a). In our original system (v1), we fused DNMT3L, a catalytically inactive accessory protein that binds to DNMT3A^18,19^ to rTetR, such that adding doxycycline (dox) to the cells recruited rTetR-DNMT3L to the citrine promoter. Methylation of the citrine promoter by endogenous DNMT3A then silenced citrine expression, which could be used as a proxy for DNMT3A activity. After introducing a library of single-guide RNAs (sgRNAs) to the cells containing this reporter system, differential silencing upon addition of dox was used to sort cells based on the activity of the DNMT3A mutants resulting from each sgRNA (Figure 1b).

**Figure 1:**
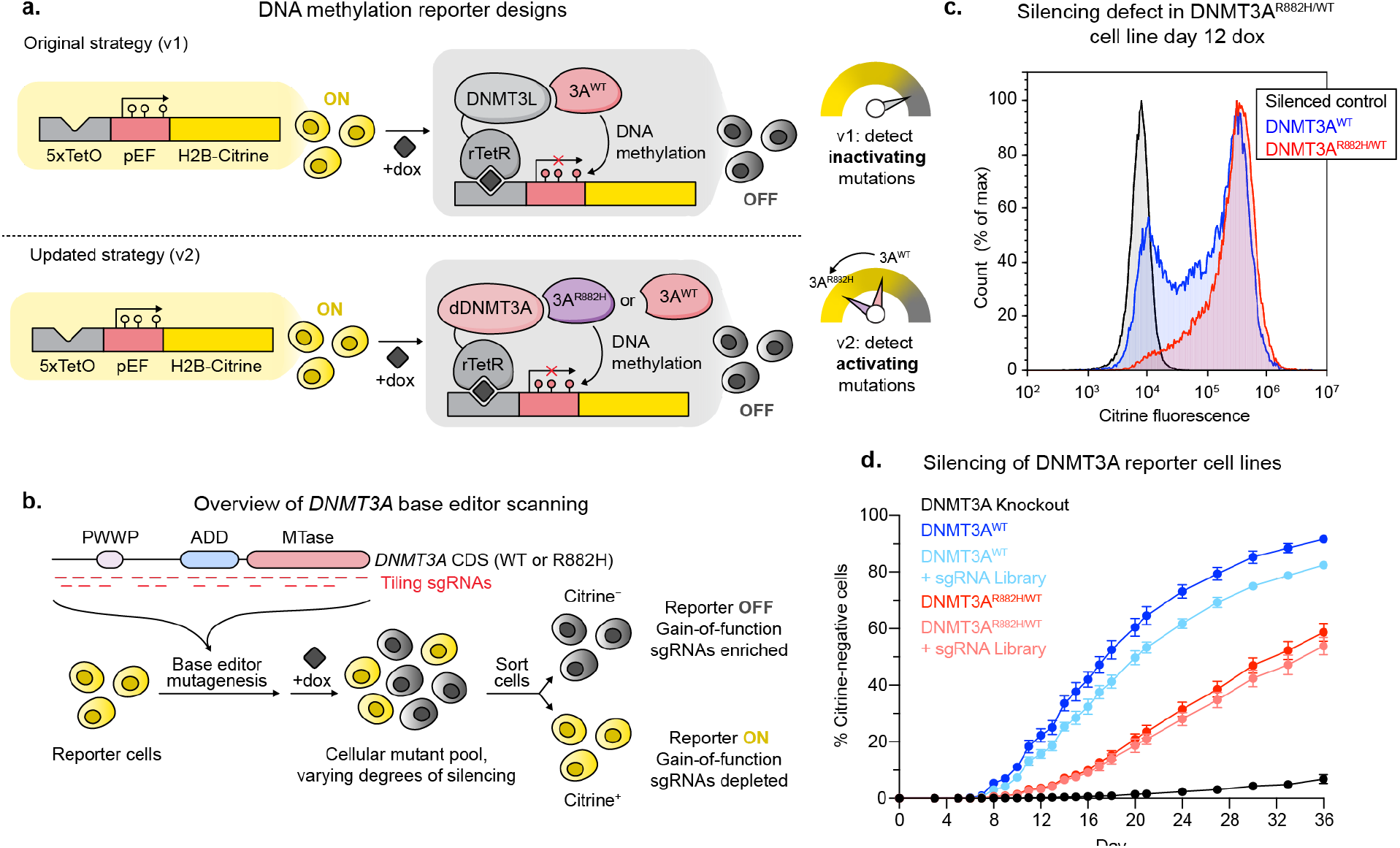
Strategy for and validation of DNMT3A activating screen. a) Cartoon showing the original base editor scanning and updated strategy for identifying *DNMT3A* activating mutations. By lowering the background activity of our reporter, we decreased the probability that activating mutations would be hidden by the limited dynamic range between baseline and maximum DNMT3A activity. dDNMT3A = codon-switched DNMT3A2^E756A^. b) Cartoon showing our strategy for base editor scanning and separating cells based on DNMT3A activity. c) Histogram showing silencing of citrine reporter in cells with DNMT3A^WT^ or NMT3A^WT/R882H^ after 12 days of dox treatment. d) Line plot showing silencing over time of DNMT3A^WT^ or DNMT3A^WT/R882H^ cells with and without the base editing library. Data are mean ± SD of n = 3 replicates.

While this reporter setup successfully identified inactivating *DNMT3A* mutations, it was not optimized to identify activating mutations since DNMT3L itself activates DNMT3A,^18^ resulting in a much higher baseline level of silencing (Figure 1a). Therefore, we modified the reporter system to address these limitations. First, we swapped DNMT3L for a catalytically dead version of DNMT3A (dDNMT3A) harboring a mutation to the catalytic glutamate (E756A),^20^ which can also recruit endogenous DNMT3A due to DNMT3A’s active state being a tetramer.^2^ To prevent base editing of dDNMT3A, we adjusted the codons encoding this copy such that this recruiting copy would not be targeted by our sgRNA library (see Methods). To further decrease the background silencing rate of our reporter system and also identify base edits that could activate the most common clinical DNMT3A mutant, we also knocked in a heterozygous DNMT3A^R882H^ mutation into our K562-based reporter cell line, resulting in isogenic DNMT3A^WT/WT/WT^ and DNMT3A^WT/R882H/R882H^ cell lines (Figure S1). We then performed parallel screens in these two cell lines (termed DNMT3A^WT^ and DNMT3A^WT/R882H^, respectively). Altogether, by using dDNMT3A as a recruitment bait and introducing the R882H mutation, we aimed to create a larger differential between the reporter silencing rate in cells that receive an activating mutation versus a neutral mutation (Figure 1a).

As expected, the DNMT3A^WT/R882H^ cell line silenced the citrine reporter more slowly than DNMT3A^WT^ cells (Figure 1c). In order to verify that citrine silencing reported on DNMT3A activity, we knocked out either *DNMT3A* or the other human *de novo* DNA methyltransferase *DNMT3B* in our reporter cell lines, treated the cells with dox, and tracked the silencing of the reporter over time, confirming that reporter silencing is DNMT3A-dependent (Figure S2a). A silencing time course in each of our cell lines with or without the sgRNA library demonstrated that the DNMT3A^WT/R882H^ cell line silences more slowly than the DNMT3A^WT^ cell line across the entire time course of the experiment (Figure 1d).

### Paired base editor screens identify activating mutations

To conduct DNMT3A base editor scanning, we next transduced a lentiviral library containing our previously employed DNMT3A sgRNA library, which contains 730 sgRNAs and the C-to-T base editor BE3.9max^15^, into the reporter cell lines. sgRNAs were annotated to edit within a given window from the +4 to +8 positions of the sgRNA with the protospacer adjacent motif (PAM) defined as positions +21–23. We treated transduced cells with dox to recruit rTetR-dDNMT3A and subsequently endogenous DNMT3A to the reporter and then sorted for cells that had silenced their citrine reporter genes, using fully silenced cells as a gating control (Figure S2b). Sorted cells were collected when the cells were ∼10-15% silenced, corresponding to day 10 for DNMT3A^WT^ or day 17 for DNMT3A^WT/R882H^. By comparing the abundance of cells harboring a particular sgRNA between the citrine negative (dark) and unsorted cells, we identified sgRNAs that accelerate citrine reporter silencing by base editing *DNMT3A* (Figure 2a,b).

**Figure 2:**
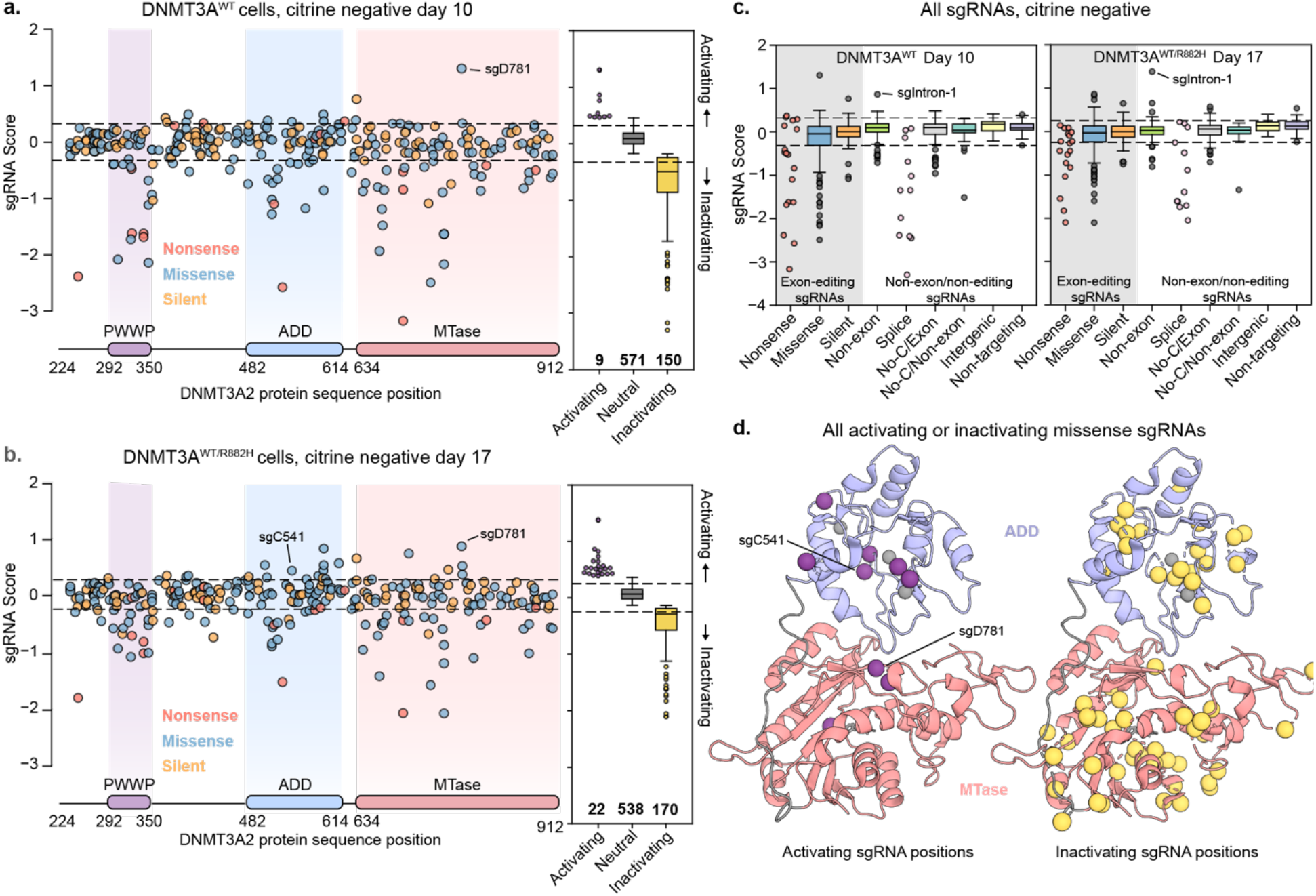
Results of base editor screens in paired isogenic cell lines. a) Scatter plot of sgRNA enrichment across the coding sequence of *DNMT3A* in DNMT3A^WT^ cells. sgRNA score corresponds to log_2_ fold-change of citrine-negative cells/unsorted cells. b) Same as a) but in DNMT3A^WT/R882H^ cells. c) Box plot showing enrichment of sgRNAs sorted by category in both screens. sgRNAs in the gray box are predicted to edit within a *DNMT3A* exon, while those in other categories either edit outside of *DNMT3A* exons or are not predicted to edit. d) Pymol cartoon model showing predicted mutated residues from missense activating (left) and inactivating (right) sgRNAs mapped to the crystal structure of the DNMT3A ADD and MTase domains (PDB: 4u7p).^10^

Comparison of the parallel screens revealed that many more sgRNAs scored as activating in the DNMT3A^WT/R882H^ cell line than in the DNMT3A^WT^ cell line (Figure 2a,b), supporting the notion that lowering DNMT3A background activity increases the dynamic range for identifying activating mutants. Despite these differences, sgRNAs predicted to introduce nonsense mutations were depleted in both cell lines and control sgRNAs were largely unchanged between sorted and unsorted cells (Figure 2c), indicating that the base editor scans faithfully reported on DNMT3A activity. In both cell lines, activating sgRNAs enriched in the ADD and MTase domains, suggesting that they may disrupt autoinhibitory functions of the ADD domain (Figure 2d). Conversely, inactivating sgRNAs generally spanned the entire protein, as we observed previously^12^ (Figure 2d).

Interestingly, sgRNAs predicted to introduce nonsense and splice site disruption mutations exhibited a greater negative effect in the DNMT3A^WT^ cell line than in the DNMT3A^WT/R882H^ cell line (Figure 2c), perhaps due to knockout of the dominant negative R882H copy being advantageous for activity. Of note, one sgRNA annotated as intronic (sgIntron-1, editing window chr2:25243984-25243988) was enriched in both cell lines. Out-of-window editing by sgIntron-1 could potentially edit the exonic region of DNMT3A, so we highlighted it for future investigation along with the enriched sgRNAs predicted to introduce missense mutations.

### Cellular and *in vitro* characterization of activating mutations

To directly compare the base editor results, we plotted the enrichment score of each sgRNA in each cell line against each other (Figure 3a). While more activating mutations were identified in our DNMT3A^WT/R882H^ screen than in the DNMT3A^WT^ cell line, the sgRNA scores were still correlated across each cell line. Next, we sought to validate top activating sgRNAs by introducing them into reporter cells individually. First, we confirmed that DNMT3A base editing did not significantly destabilize the protein (Figure S3). Next, we tested the silencing ability of the base-edited cells compared to cells containing a control nontargeting (Luc) sgRNA (Figure 3b). In this format, five of the seven tested sgRNAs were validated. Of note, while sgR631 was not fully validated, low levels of activation were observed in one replicate of this experiment (Figure S4a,b), suggesting that variability in editing efficiency might sometimes obscure our ability to detect DNMT3A activation.

**Figure 3:**
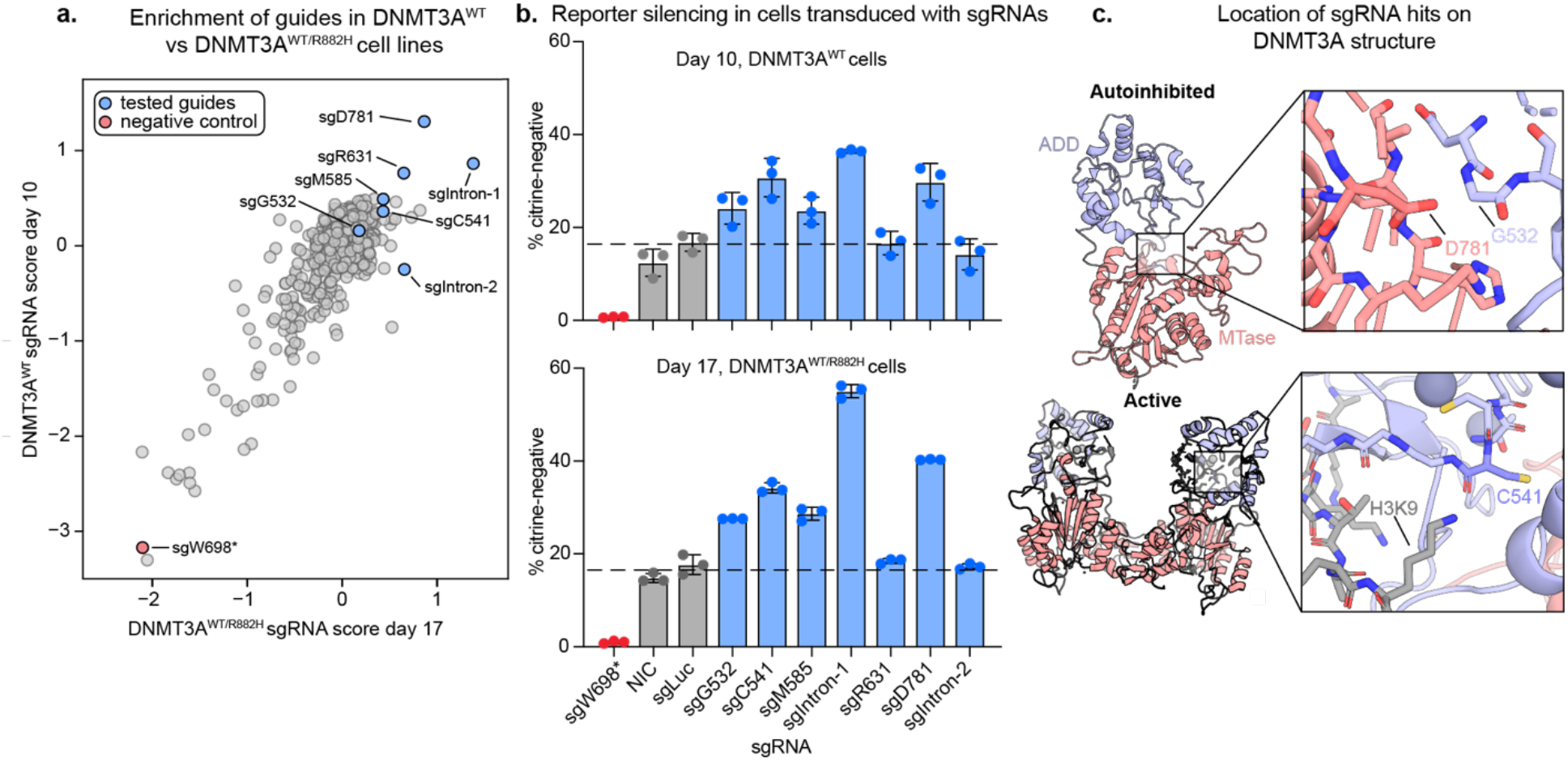
Validation of sgRNA hits. a) Scatter plot showing sgRNA scores in DNMT3A^WT^ vs DNMT3A^WT/R882H^ reporter cells. sgRNAs tested in b) and c) are highlighted. Spearman R = 0.51, p=8.1×10^−44^ b) Bar plots showing % silenced cells treated with single sgRNAs after 10 (DNMT3A^WT^) or 17 (DNMT3A^WT/R882H^) days of dox treatment. sgLuc = Luciferase-targeting sgRNA, NIC = non-infection control. Data are mean ± SD of n = 3 replicates. See figure S4 for additional sgRNAs and replicates. c) PyMOL models showing the structural location of key activating mutants. PDB IDs: 4u7t (active) and 4u7p (autoinhibited).^10^

After performing silencing assays on cells transduced with individual sgRNAs, we next genotyped the edited cells and compared the observed mutations with the predicted edits. Of the validated sgRNAs, sgD781 introduced the predicted D781N mutation (Figure 4a), which is positioned directly across the autoinhibitory interface from the activating mutation found in our previous work, G532N^12^ (Figure 3c). sgC541 introduced the predicted C541Y mutation in the ADD domain, which lies close to the interaction site for K9 of the activating H3K4me0 peptide. Interestingly, this mutation is distal to the autoinhibitory interface that is perturbed by the G532N or D781N mutants.

**Figure 4:**
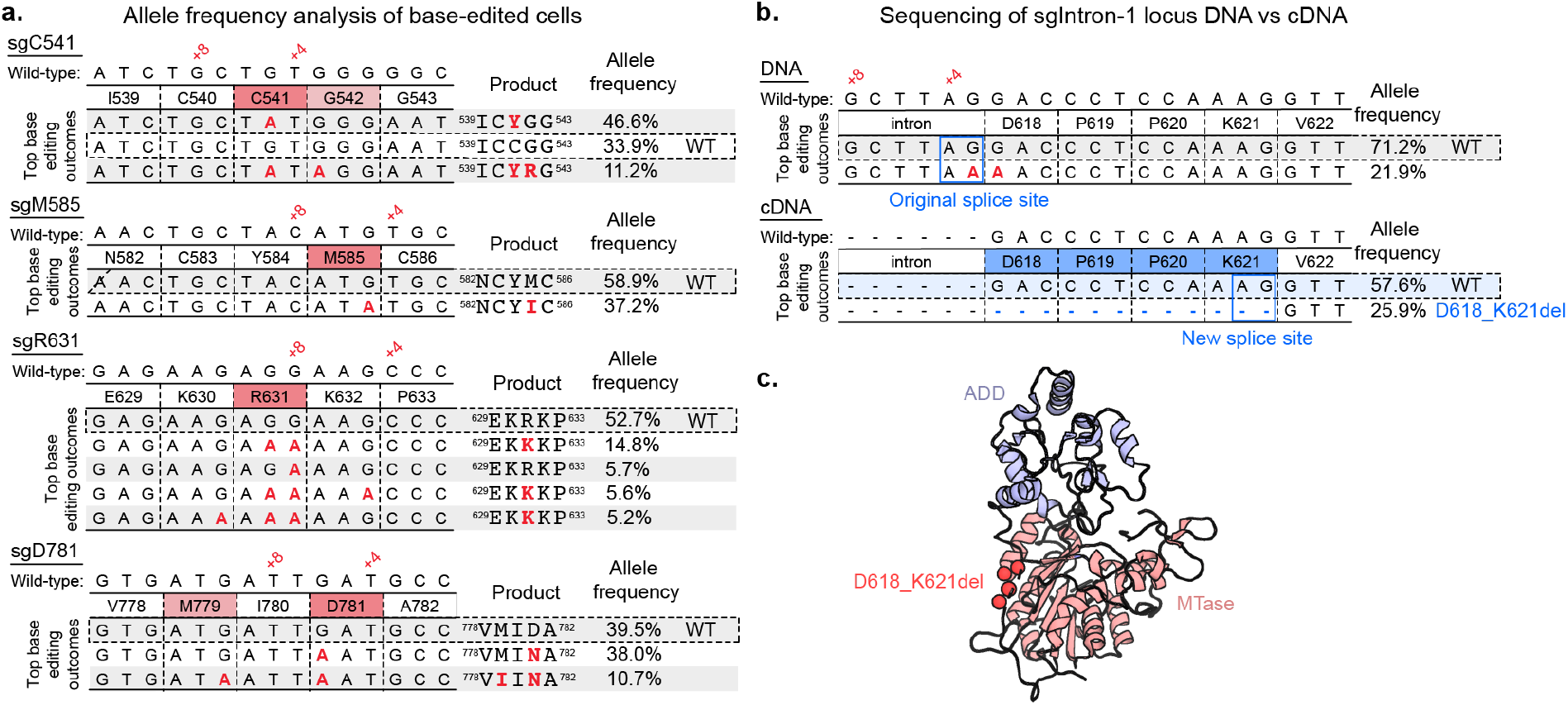
Genotyping of sgRNA hits. a) Tables showing base editing outcomes with >5% allele frequency from treatment of DNMT3A^WT^ reporter cells with single missense sgRNAs (See Figure S5 for DNMT3A^WT/R882H^ cell line). Analysis performed with CRISPResso2^21^. b) Scheme showing genomic DNA and cDNA sequencing of DNMT3A^WT^ reporter cells treated with sgIntron-1. (See Figure S5 for DNMT3A^WT/R882H^ cell line). c) PyMOL model showing location of D618_K621del (PDB ID 4u7p).

sgIntron1 was strongly activating as well despite its editing window being in an intron (Figure 3b). Genotyping of the targeted genomic locus revealed editing at the +3 position of the sgRNA, one base pair outside of the canonical editing window (Figure 4b). Notably, the edited C complements the G of a splice acceptor site. Therefore, we hypothesized that the editing of this residue could induce alternative splicing that utilizes an “AG” sequence motif 12 base pairs into the next exon, causing a four-amino acid deletion in the resulting translated protein (D618_K621del). To test this hypothesis, we sequenced the *DNMT3A* cDNA from cells transduced with sgIntron-1 and observed that 29% and 38% of transcripts from DNMT3A^WT^ and DNMT3A^WT/R882H^ cells, respectively, were alternatively spliced as predicted (Figure 4b). The resulting translation contains a deletion of residues 618-621 in the disordered linker region between the ADD and MTase domains (Figure 4c). In addition, the cryptic splice site leading to the D618_K621del mutation is also found in several species of colobinae monkeys, indicating that this mutation has been sampled in the course of DNMT3A evolution (Figure S6). This demonstrates an unexpected mechanism for generating a deletion mutation using a base editor. Overall, we identified the activating mutants D781N, C541Y, and D618_K621del as strongly activating in our reporter assay and selected them to study *in vitro*.

Having validated a subset of activating sgRNAs in the cellular reporter system, we next evaluated the activity of the corresponding purified DNMT3A mutants compared to WT DNMT3A and the single activating hit from our previous work^12^, G532N. In a biochemical methyltransferase assay, G532N demonstrated ∼9-fold higher activity than WT DNMT3A, while the mutants identified in the updated screen were ∼4 fold more active than WT (Figure 5a). We also confirmed that our mutants are stable by differential scanning fluorimetry (Table S3). We next considered whether these activating mutants are stimulated by H3K4me0. Interestingly, all mutants showed increased enzymatic activity in the presence of 5 µM H3K4me0 peptide (K_D_ for H3K4me0 binding to DNMT3A ADD domain is 1.75 µM ± 0.11^10^) (Figure 5b). These results, along with the positions of the D618_K621del and C541Y mutations in the protein structure, suggest that the mutations do not activate the protein by fully relieving the known autoinhibition mechanism, although they do suggest that these mutants could still be perturbing the equilibrium between the autoinhibited and active conformations. Overall, we identified three mutations to DNMT3A through base editor screens that are in and around the ADD domain and are activating both in our cellular reporter system and *in vitro*.

**Figure 5:**
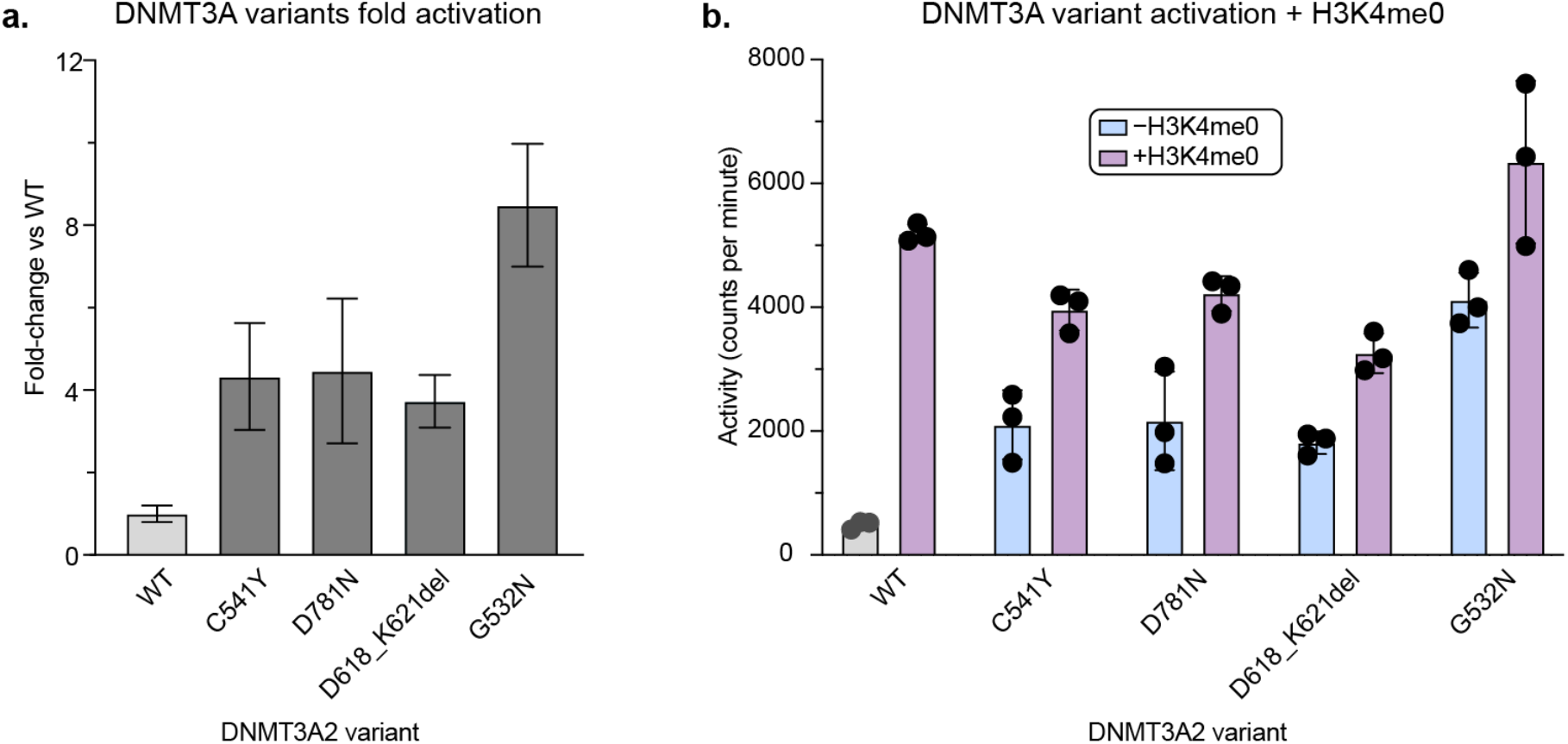
Biochemical validation of activating mutants. a) Bar plot showing fold-change in DNMT3A methyltransferase activity of mutants vs WT, based on data in b). Fold-change errors were propagated from the individual standard deviations. b) Bar plot showing methyltransferase activity of DNMT3A mutants ± 5 µM H3K4me0. Data are mean ± SD of n = 3 replicates. See Figure S7 for SDS-PAGE gel of purified proteins and replicate 2.

## Discussion

By innovating new DNMT3A reporter strategies and enzyme hypomorphs with base editing technology, we expanded our ability to discover activating mutations to DNMT3A. Our approach demonstrates the ability of the adapted DNMT3A reporter strategy to identify new activating mutants with more intermediate effects, which was not possible with our initial strategy. These results highlight the importance of considering the baseline activity of the protein of interest and selection in designing an appropriate screening strategy. Furthermore, we provide a blueprint for how parallel base editor screens can be performed in isogenic cell lines. While the mutant DNMT3A^WT/R882H^ cell line provided additional dynamic range in which to detect activating mutations, ultimately all activating mutations identified in the mutant cell line were also activating in the WT cell line. The ability of these mutants to increase DNMT3A activity in both the WT and the most common clinical mutant contexts further highlights these areas of DNMT3A as valuable and broadly applicable pharmaceutical targets. We anticipate that base editing approaches in isogenic cell lines could be a valuable strategy to investigate other proteins (i.e., oncogenes, tumor suppressors) and their corresponding mutants of interest. We also highlight how well-designed selection strategies can enable the identification of hypermorphic variants of a protein of interest with base editor scanning.

One limitation of this study was the use of a cytosine base editor with an “NGG” protospacer-adjacent motif requirement, which reduced the number of accessible mutations. Despite these limitations, our optimized screening strategy successfully identified several new activating mutations, suggesting that future studies employing next-generation SpG and SpRY cytidine and adenosine base editors^22^ could uncover even more activating mutations. Furthermore, the ability of base editing to target splice sites—which in the past has been used to disrupt genes,^23^ induce exon skipping,^24^ or as a control for LOF mutations—could serve as an interesting tool to probe splice sites in clinically-relevant proteins for potential gain-of-function (GOF) mutations caused by alternative splicing. Altogether, our results suggest that base editing of splice sites might be a good strategy to explore cryptic splicing variants of different proteins of interest.

In this study, we identified and validated three new mutations that demonstrate novel ways to activate DNMT3A, which point to the ADD domain as a master regulator of DNMT3A function and a possible target site to potentiate DNMT3A activity. The D781N mutation likely demonstrates another example of disrupting the ADD–MTase autoinhibitory interface. The D618_K621del activating mutation alters a disordered linker region of DNMT3A, nominating this unstructured region as potentially important for DNMT3A regulation. The location of C541Y on the structure of the protein indicates that areas of the ADD domain beyond the known autoinhibitory interface can also be involved in DNMT3A activation. Notably, the C541Y mutation involves a cysteine to tyrosine substitution, suggesting that covalent modification of the cysteine by a small molecule electrophile might phenocopy the effect.^25^ Given the large body of evidence indicating that DNMT3A LOF promotes developmental disorders and leukemogenesis in hematopoietic disorders development of small molecules that could activate DNMT3A activity would be of significant interest. Exploiting these activating mutations could be critically instructive in identifying functional sites for small molecule agonists of DNMT3A. We anticipate that combining innovative selection strategies with base editing mutagenesis could eventually help inform drug discovery efforts to recover activity in other tumor suppressor targets.

## Methods

### Safety statement

No unexpected or unusually high safety hazards were encountered.

### General plasmid construction

sgRNA plasmids were cloned as described in ref ^12^. Gibson Assembly with NEBuilder HiFi (New England Biolabs) was used to clone additional plasmids. NEB Stable (lentiviral vectors) and NEB 5-alpha (other plasmids) (New England Biolabs) were used as cloning strains. Final constructs were validated by Sanger (Quintara Biosciences) or Oxford Nanopore (Plasmidsaurus) sequencing. Plasmids and oligonucleotides are provided in Tables S1 and S2, respectively.

### Construction of rTetR-dDNMT3A reporter

To recruit endogenous DNMT3A to the reporter promoter, a catalytically dead DNMT3A2 mutant (DNMT3A2^E756A^) was fused to rTetR, in an adaptation of our previously reported strategy^12^. The gene encoding the recruiting DNMT3A2^E756A^ was engineered to prevent its recognition and mutagenesis by the base editor scanning DNMT3A2 sgRNA library. A custom python script was used to insert synonymous substitutions *in silico* throughout the DNMT3A2 coding sequence to block sgRNA recognition. Briefly, for each library sgRNA recognizing an exon or exon boundary, the ‘NGG’ protospacer-adjacent motif (PAM) was disrupted where possible; otherwise, protospacer positions were mutated to create at least 2 mismatches, beginning at the PAM-proximal and moving toward the PAM-distal end. After considering all sgRNAs, protospacer mismatches were enumerated. For any sgRNA with 3 or fewer protospacer mutations, none of which were in the 6 most PAM-proximal positions, an additional mismatch was introduced if possible. This resulted in a gene encoding wild-type DNMT3A2 but containing mismatches with each library sgRNA. Finally, the E756A substitution was introduced. The DNMT3A2^E756A^ gene was synthesized by IDT and cloned into pRRL-pEF-H2B-mCherry-T2A-rTetR-Dest-SV40 (Addgene #186968).

### Cell culture

K562 (ATCC), 293FT (Thermo Fisher Scientific), and HEK293T (a gift from B. E. Bernstein, Massachusetts General Hospital) cell lines were used in this study. All cell lines were cultured at 37 °C and 5% CO2 in a humidified incubator and were tested for mycoplasma. All media were supplemented with 100 U ml^−1^ penicillin, 100 μg ml^−1^ streptomycin (Gibco), and 10% FBS (Peak Serum). K562s were cultured in RPMI-1640 (Gibco), HEK293Ts were cultured in DMEM (Gibco), and 293FTs were cultured in DMEM with 2 mM GlutaMAX (Gibco) and 1× MEM nonessential amino acids (Gibco).

### Lentiviral transduction

To produce lentivirus, GAG/POL and VSVG plasmids were co-transfected with transfer plasmid into HEK293Ts or HEK293FTs using FuGENE HD (Promega, 3.33:1 FuGENE:DNA), or lipofectamine 3000 Reagent (Thermo Fisher Scientific). Medium was then replaced after 6– 8 hours. 48–60 hours later, the medium was passed through a 0.45-μm filter and either used immediately or snap-frozen for storage at −80 °C. To generate reporter cell lines and introduce sgRNAs, K562s were transduced with the appropriate lentivirus by spinfection (1,800g, 90 min, 37 °C) with 12 μg ml^−1^ polybrene (Santa Cruz Biotechnology). 2 μg ml^−1^ puromycin (Gibco) or fluorescence-activated cell sorting (FACS) were used to select for transduced cells. A clonal methylation reporter cell line (DNMT3A^WT^) was first derived and used to generate the DNMT3A^WT/R882H^ cell line. This clonal cell line or the DNMT3A^WT/R882H^ clonal cell line was used for all individual sgRNA transductions.

### Generation of DNMT3A^WT/R882H^ reporter cell line

All CRISPR reagents were obtained from Integrated DNA Technologies (IDT) unless otherwise specified. In order to knock in the R882H mutation to DNMT3A in our reporter cell line, CRISPOR^26^ was used to design an appropriate guide RNA targeting the DNMT3A R882 locus (see guide sequences in Table S2). The guide RNA was ordered as an Alt-R CRISPR-Cas9 crRNA. The Alt-R HDR Donor Oligo was designed using the IDT web tool. To prepare the gRNA complex, crRNA and Alt-R CRISPR-Cas9 tracrRNA were mixed in equimolar ratio, heated to 95C for 5 min, then allowed to cool to room temperature. To prepare the RNP complex, the gRNA complex was combined with Alt-R Cas9 enzyme and incubated at RT for 20 min. The Neon™ Transfection System was used to electroporate 2 ×10^5^ DNMT3A^WT^ reporter cells with the assembled RNP complex and HDR Donor. Electroporated cells were recovered in antibiotic-free media for 72 hours. Single cell clones were sorted by FACS and screened by amplicon sanger sequencing for the presence of the R882H mutation. The final clone was confirmed to have two copies or the DNMT3A^R882H^ allele and one copy of the DNMT3A^WT^ allele by Illumina amplicon sequencing (Figure S1).

### Reporter silencing assays

Reporter silencing assays were conducted as in ref ^12^ with the following changes: cells were treated with 1ug ml^−1^ dox (Sigma-Aldrich), and gates were set based on reference reporter cells cultured in parallel without dox and reference fully silenced reporter cells (See gating scheme in Figure S2).

### Genotyping

The QIAamp DNA Blood Mini kit (Qiagen) was used to purify genomic DNA. 100 ng of DNA was inputted into a first round of PCR (27 cycles, Q5 hot start high-fidelity DNA polymerase (New England Biolabs)) to attach common overhangs and amplify the target locus. To attach barcoded adapters, 1 µL of each PCR1 reaction was amplified in a second round of PCR (8 cycles). Primer sequences are provided in Table S2. Gel extraction (Zymo) was used to purify final amplicons, which were then sequenced on an Illumina MiSeq. Data were processed using CRISPResso2^21^.

### Reverse transcription and sequencing of sgIntron_1 locus

The RNeasy Mini Kit (Qiagen) was used to isolate RNA. Then, MultiScribe Reverse Transcriptase (Invitrogen) was used with 10x RT Random Primers (Invitrogen) to amplify 1.8 ug of RNA for each sample. ∼50 ng of the resulting cDNA was then used as template for amplification with pEG1.136_1R and pEG1.236_1F for 40 cycles of PCR. The resulting products were then barcoded and sequenced in the same manner as our DNA genotyping.

### Immunoblotting

Immunoblotting was performed as in ref XX. The following antibodies and dilutions were used: Primary antibodies: DNMT3A (Cell Signaling Technology, catalog no. 32578, D2H4B, 1:5,000), GAPDH (Santa Cruz Biotechnology, catalog no. sc-47724, 0411, 1:8,000). Secondary antibodies: anti-rabbit IgG HRP conjugate (Promega, catalog no. W4011, 1:100,000 for DNMT3A), anti-mouse IgG HRP conjugate (Promega, catalog no. W4021, 1:400,000 for GAPDH). Immunoblots were visualized using SuperSignal West Femto (DNMT3A) or SuperSignal West Pico PLUS (GAPDH) chemiluminescent substrates (Thermo Fisher Scientific).

### Base editor scanning

This study employs the sgRNA library described in ref ^12^. Titering of library lentivirus was performed by measuring cell counts of transduced cells after puromycin selection. 30 × 10^6^ reporter cells per cell line were transduced with library lentivirus at a multiplicity of infection <0.3 and selected with puromycin for 7 days. Following selection and expansion cells were split into three replicates and treated with 1 ug/mL dox. DNMT3A^WT^ reporter cells were passaged on days 3, 6, and 7 of dox treatment, and a portion of the cells were sorted on day 10 for sequencing analysis. DNMT3A^WT/R882H^ reporter cells were passaged on days 3, 6, 9, 11, 14, and 17 of dox treatment, and a portion of the cells were sorted on day 17 for sequencing analysis. Cells were sorted on a FACSAria II (BD), collecting citrine+, citrine− and unsorted (all mCherry+) cells. Fully silenced control cells were used for citrine gating (see Figure S2). Library preparation for Illumina sequencing and data analysis was performed as in ref ^12^. Library sgRNA annotations and base editor scanning data are provided in Tables S4 and S5.

### Protein expression and purification

Full-length DNMT3A2 was recombinantly expressed and purified according to ref. ^12^ with the following modifications: Base buffer consisted of 50 mM Tris-HCl, pH 8.0 cold, 500 mM NaCl, 1 mM TCEP, and lysis buffer was supplemented with 10 mM imidazole and 0.2% Triton X-100. DNMT3A storage buffer consisted of Base buffer but with 300mM NaCl.

### Activity assays

DNMT3A activity was measured using the protocol described in ref ^12^. For histone stimulation experiments, 5 μM peptide was added prior to addition of enzyme. The following peptides were used: H3K4me0 (ARTKQTARKSTG-NH2, Biomatik), H3K36me2 (histone H3 (21–44)-GK (biotin), Anaspec, catalog nos. AS-64440 and AS-64442).

### Differential scanning fluorimetry

First, 2 μM purified DNMT3A2 was incubated in triplicate with 5× SYPRO Orange (Thermo Fisher Scientific) in assay buffer (50 mM Tris-HCl, pH 8, 300 mM NaCl, 1mM TCEP). Samples were heated from 10 °C to 95 °C (10 s at each 0.5 °C step) using a CFX Connect qPCR with CFX Manager software (Bio-Rad). Melting temperatures were calculated with DSFWorld^27^ (by model 2).

### Colobinae monkey sequence analysis

The jackhmmer webserver^28^ (accessed April 2020) was used with human DNMT3A (UniProt: Q9Y6K1) as the query sequence and an E-value cutoff of 0.01 to generate a multiple sequence alignment. This alignment was then manually searched for sequences containing the 4 amino acid deletion generated by sgIntron-1, and the resulting sequences were realigned using Geneious. Sequence identities were also calculated using Geneious.

## Supporting information

WT cell line screen results

WT/R882H cell line results

## Acknowledgements

We would like to thank members of the Liau Lab and others for insightful comments and discussion on the manuscript, in particular J. Morris, H. S. Kwok, and S. Berry. We additionally thank Z. Niziolek and C. Maesner at the Bauer Core Facility at Harvard University for help with FACS, and D. Bolduc for biochemistry advice.

E.M.G. and N.Z.L. were supported by National Science Foundation Graduate Research Fellowships (grant no. DGE1745303). E.M.G. was also supported by a Landry Cancer Biology Fellowship. B.B.L. is a Damon Runyon-Rachleff Innovator supported in part by the Damon Runyon Cancer Research Foundation (grant no. DDR 60S-20). This research was additionally supported by award no. 1DP2GM137494 from the National Institute of General Medical Sciences and startup funds from Harvard University.

