## Supplemental Information for "Base Editor Scanning Reveals Activating Mutations of DNMT3A"

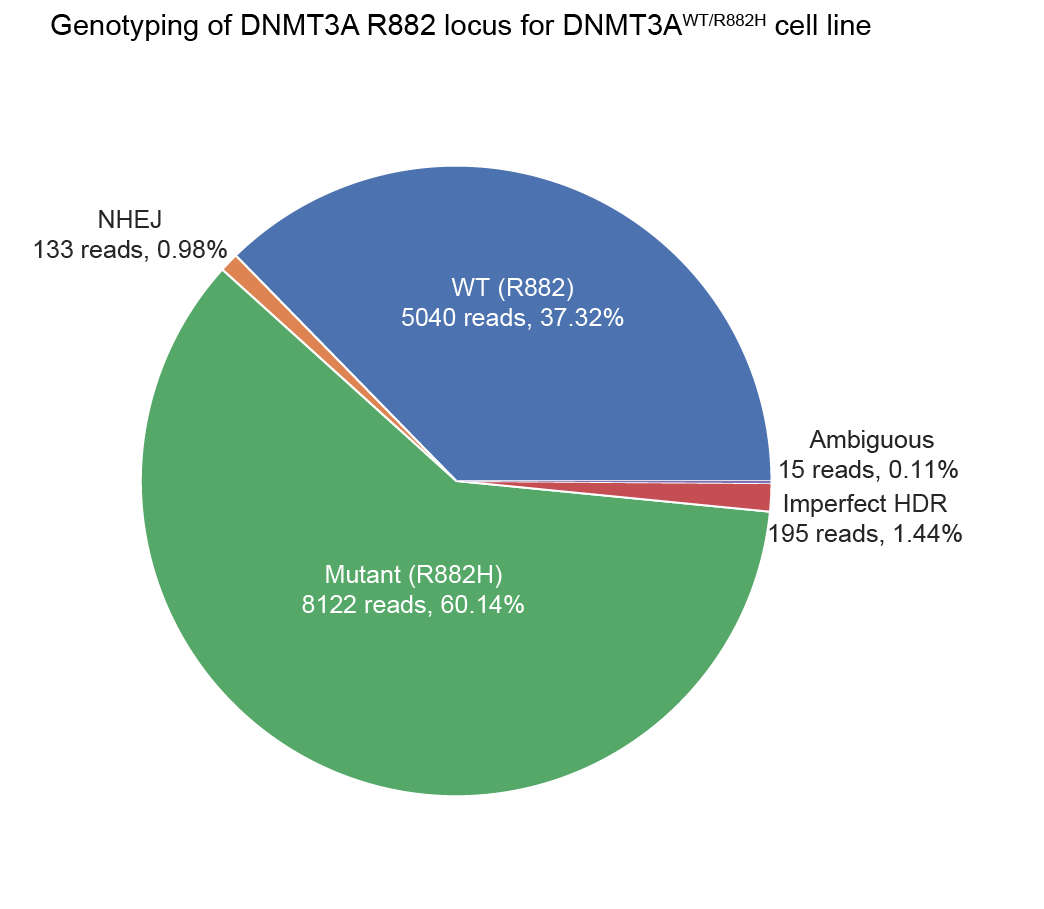

**Figure S1: Pie chart showing targeted deep sequencing of R882 codon of *DNMT3A* in DNMT3A^WT/R882H^ cells.** K562 cells are triploid, and these genotyping results are consistent with our cell line containing 2 copies of DNMT3A^R882H^ and one copy of DNMT3A^WT^. Plot generated with CRISPResso2

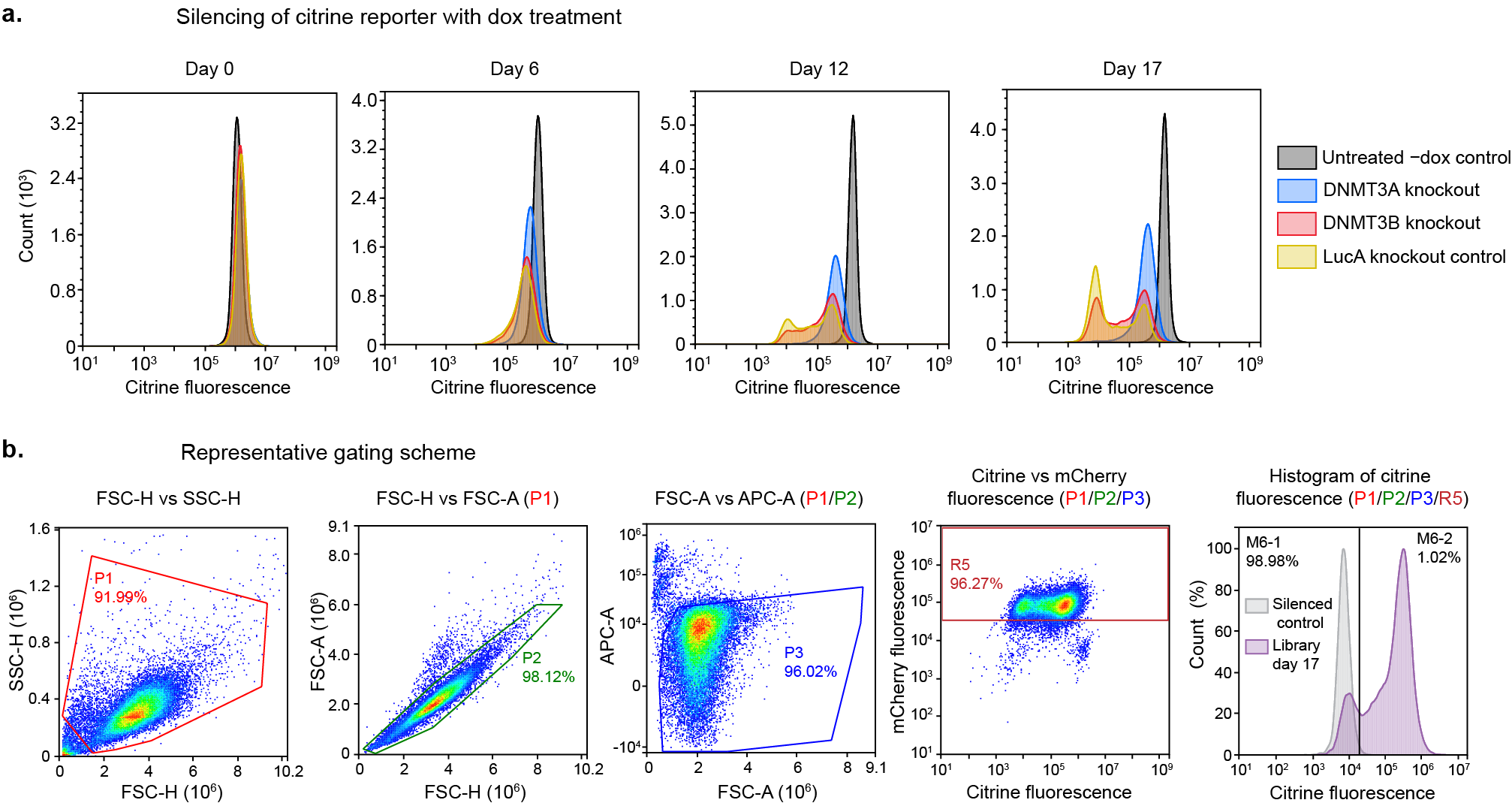

**Figure S2: Silencing dynamics of dDNMT3A reporter system and representative gating scheme.**

a.) Histograms of DNMT3A^WT^ reporter cells treated with DNMT3A knockout, DNMT3B knockout, or LucA knockout (control) sgRNAs and Cas9 after 0, 6, 12, and 17 days of 1 ug/mL doxycycline treatment.
b.) Representative gating scheme for sorting cells in our screen. Events were first gated for morphologically typical K562 cells, then for events corresponding to single cells. Then, helix-IR staining was used to gate out dead cells. Finally, the population corresponding to healthy, single cells was gated for mCherry+ cells and assessed for citrine fluorescence. Cells were sorted based on a citrine gate where ~1% of fully silenced control cells are citrine+.

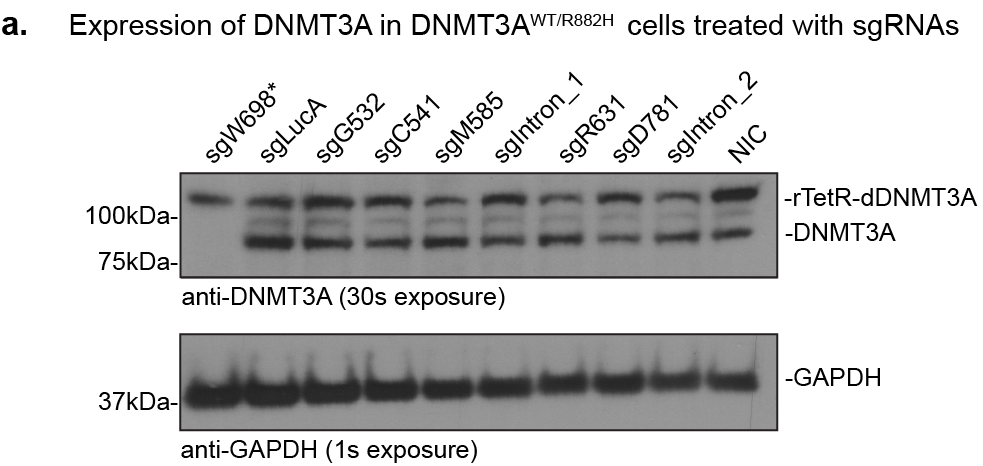

**Figure S3: Western Blot of reporter cells transduced with base editing sgRNAs**. sgW698* causes depletion of endogenous DNMT3A levels but not rTetR-dDNMT3A levels. Other sgRNAs tested (the same set as used in Figure 3b) do not dramatically affect protein expression levels.

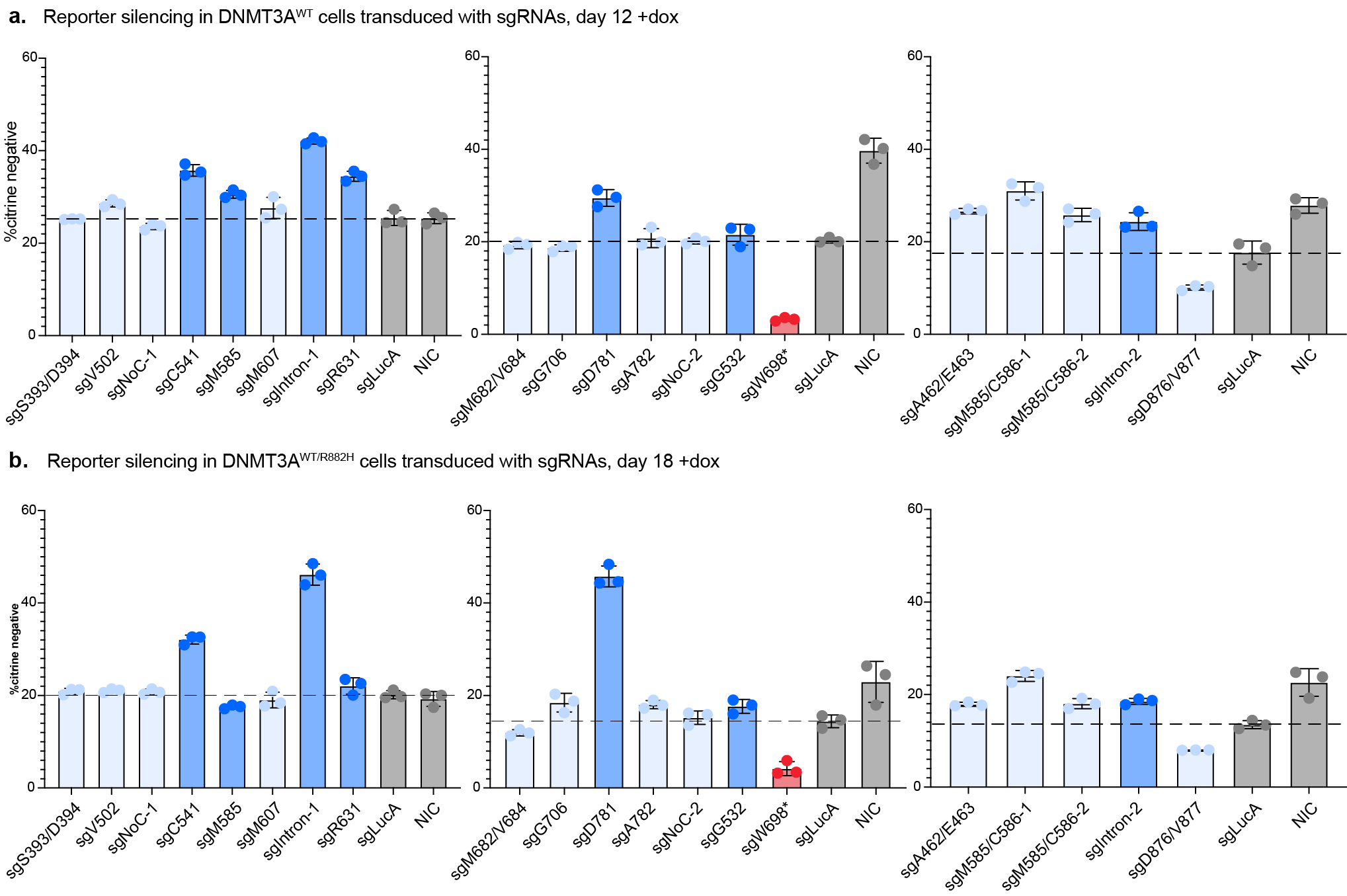

**Figure S4: Additional sgRNAs tested.**

a. Bar plots showing % silenced DNMT3A^WT^ cells treated with sgRNAs after 12 days of dox treatment. LucA = Luciferase targeting sgRNA, NIC = Non-infection control. Dark blue, red and gray bars indicate biological replicates for guides shown in Figure 3b, and light blue bars indicate one biological replicate in technical triplicate for additional sgRNAs. Data are mean ± SD of n = 3 replicates.
b. Bar plots showing % silenced DNMT3A^WT/R882H^ cells after 17 days of dox treatment. Data are mean ± SD of n = 3 replicates.

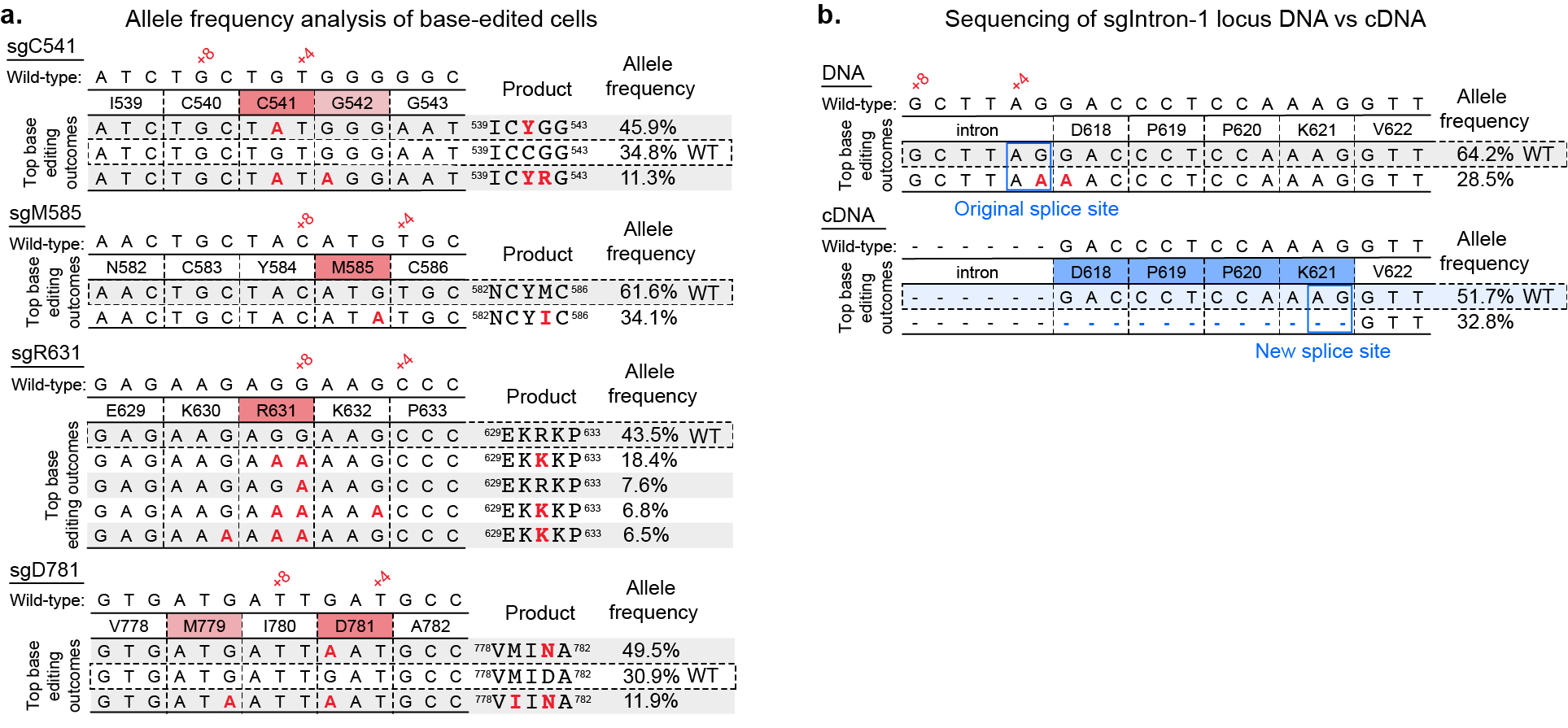

**Figure S5: Genotyping of DNMT3A^WT/R882H^ cells treated with sgRNAs.**

a. Tables showing base editing outcomes with >5% allele frequency from treatment of DNMT3A^WT/R882H^ reporter cells with single missense sgRNAs.
b. Scheme showing genomic DNA and transcript cDNA sequencing of cells treated with sgIntron-1.

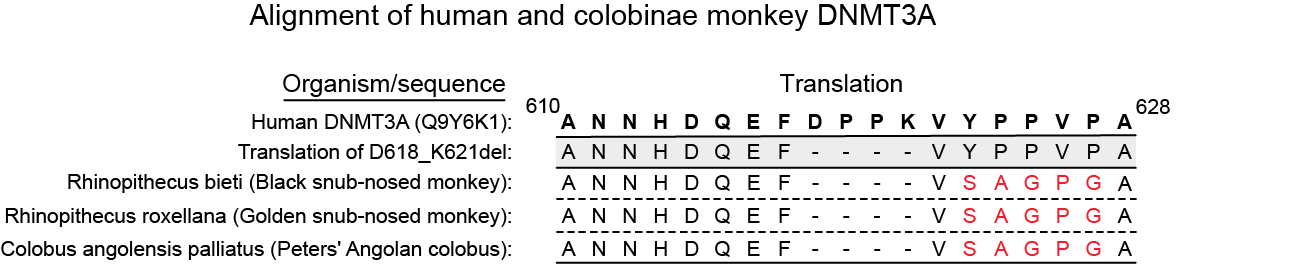

Overall sequence identity

|  | **Human** | **Black snub-nosed** | **Golden snub-nosed** | **Peters’ Angolan** |
| --- | --- | --- | --- | --- |
| **Human** | 100% | 96.8% | 96.8% | 95.3% |
| **Black snub-nosed** | 96.8% | 100% | 100% | 98.1% |
| **Golden snub-nosed** | 96.8% | 100% | 100% | 98.1% |
| **Peters’ Angolan** | 95.3% | 98.1% | 98.1% | 100% |

**Figure S6: D618_K621del mutation is found in colobinae monkeys.** Despite additional local variation in sequence between human and colobinae DNMT3A, the proteins are overall very similar. UniProt IDs: A0A2K6PNQ5_RHIRO, A0A2K6KW24_RHIBE, A0A2K5JG23_COLAP

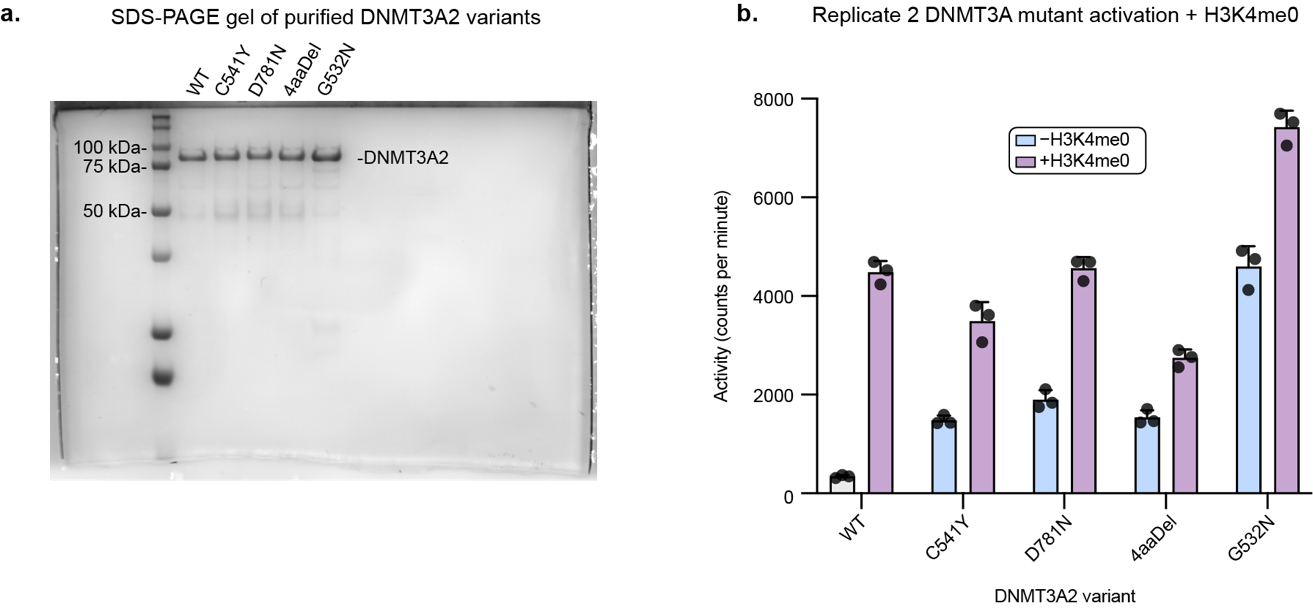

**Figure S7: Additional in vitro validation of DNMT3A variants.**

a. SDS-PAGE gel showing purified DNMT3A2 variants used in Figure 4.
b. Second replicate of activity assay shown in Figure 4b. Data are mean ± SD of n = 3 replicates

**Table S1: Plasmids constructed in this study.**

| Application | Plasmid | Description |
| --- | --- | --- |
| Base editing screen and validation | pEF-H2B-mCherry-T2A-rTetR-dDNMT3A2 | Lentiviral, mammalian expression of updated reporter |
| Bacterial Protein Expression | pET28b-His_6_-DNMT3A2-C541Y | Expression of His_6_-tagged DNMT3A2 C541Y |
|  | pET28b-His_6_-DNMT3A2-D781N | Expression of His_6_-tagged DNMT3A2 D781N |
|  | pET28b-His_6_-DNMT3A2-4aaDel | Expression of His_6_-tagged DNMT3A2 4aaDel |

**Table S2: Primers and oligos used in this study.**

| Application | Description | Sequence |
| --- | --- | --- |
| Plasmid Construction | Construction of pET28b-His_6_-DNMT3A2-D781N Forward | CTTCCTTCGCATTGATCATAACCG |
|  | Construction of pET28b-His_6_-DNMT3A2-D781N Reverse | CGGTTATGATCAATGCGAAGGAAG |
|  | Construction of pET28b-His_6_-DNMT3A2-4aaDel Forward | GGTGGATACACAAATTCTTGGTCATGG |
|  | Construction of pET28b-His_6_-DNMT3A2-4aaDel Reverse | GGTGGATACACAAATTCTTGGTCATGG |
|  | Construction of pET28b-His_6_-DNMT3A2-C541Y Forward | CCTCACGACCGCCATAGCAAATCGTAC |
|  | Construction of pET28b-His_6_-DNMT3A2-C541Y Reverse | GTACGATTTGCTATGGCGGTCGTGAGG |
|  | Construction of pEF-H2B-mCherry-T2A-rTetR-dDNMT3A2 Forward | CCTTTTTTTTGGCTCTTTGCGAACGTCGTCGCTATGGGAG |
|  | Construction of pEF-H2B-mCherry-T2A-rTetR-dDNMT3A2 Reverse | CTCCCATAGCGACGACGTTCGCAAAGAGCCAAAAAAAAGG |
| Individual sgRNAs (top= Forward, bottom= Reverse) | DNMT3A2_552 | CACCGGGCATCAATCATCACAGGGT  AAACACCCTGTGATGATTGATGCCC |
|  | DNMT3A2_414 | CACCGTCCTAAGCAGTGAGCACAAC  AAACGTTGTGCTCACTGCTTAGGAC |
|  | DNMT3A2_207 | CACCGTGTCACTCTCATCGCTGTCG  AAACCGACAGCGATGAGAGTGACAC |
|  | DNMT3A2_341 | CACCGAGCAGATGGTGCAGTAGGAC  AAACGTCCTACTGCACCATCTGCTC |
|  | DNMT3A2_617 | CACCGCATCTTATGGTGCACTGAAA  AAACTTTCAGTGCACCATAAGATGC |
|  | DNMT3A2_310 | CACCGGGTAACATTGAGGCTCCCAC  AAACGTGGGAGCCTCAATGTTACCC |
|  | DNMT3A2_396 | CACCGAAGAACATCTGGAGCCGGGA  AAACTCCCGGCTCCAGATGTTCTTC |
|  | DNMT3A2_493 | CACCGCCCCAATCACCAGATCGAAT  AAACATTCGATCTGGTGATTGGGGC |
|  | DNMT3A2_554 | CACCGTCTTTGGCATCAATCATCAC  AAACGTGATGATTGATGCCAAAGAC |
|  | DNMT3A2_381 | CACCGCGCACATGTAGCAGTTCCAG  AAACCTGGAACTGCTACATGTGCGC |
|  | DNMT3A2_422 | CACCGTGGGCTTCCTCTTCTCAGCT  AAACAGCTGAGAAGAGGAAGCCCAC |
|  | DNMT3A2_343 | CACCGCCCACAGCAGATGGTGCAGT  AAACACTGCACCATCTGCTGTGGGC |
|  | DNMT3A2_613 | CACCGTCATGAAGACAGGAAAATGC  AAACGCATTTTCCTGTCTTCATGAC |
|  | DNMT3A2_468 | CACCGCGACGTACATGATCTTCCCC  AAACGGGGAAGATCATGTACGTCGC |
|  | DNMT3A_260 | CACCGTTCTCCGCTGTGCTCTTCCG  AAACCGGAAGAGCACAGCGGAGAAC |
|  | DNMT3A_382 | CACCGCCGCACATGTAGCAGTTCCA  AAACTGGAACTGCTACATGTGCGGC |
|  | DNMT3A_384 | CACCGCCCGCACATGTAGCAGTTCC  AAACGGAACTGCTACATGTGCGGGC |
|  | DNMT3A_568 | CACCGTTTCACCAACCTGTTCATAC  AAACGTATGAACAGGTTGGTGAAAC |
| R882H Knock-in | sgRNA for CRISPR-Cas9 R882H knock-in | CCAACATGAGCCGCTTGGCG |
|  | HDR template for R882H knock-in | CAGGGTATTTGGTTTCCCAGTCCACTATACTGACGTCTCCAACATGAGCCACTTGGCGAGGCAGAGACTGCTGGGCCGGTCATGGAGCGTGCCAGTCATCC |
| Genotyping (bold=gDNA binding region) | pNL1.155 F3-sequence around R882 | ACACTCTTTCCCTACACGACGCTCTTCCGATCTNNNN**CAGCACTCACCCTGCCC** |
|  | pNL1.155 R3-sequence around R882 | TGGAGTTCAGACGTGTGCTCTTCCGATCT**TCGCTACCTCAGTTTGCCC** |
|  | pNL1.175 F1-sequence around G532 | ACACTCTTTCCCTACACGACGCTCTTCCGATCTNNNN**TCCTGGTGGTTTCTGACCCT** |
|  | pNL1.175 R1-sequence around G532 | TGGAGTTCAGACGTGTGCTCTTCCGATCT**CCAAGGTGTGCTACCTGGAA** |
|  | pEG1.103 3F-sequence exon 10 | TGGAGTTCAGACGTGTGCTCTTCCGATCT**GCCCTCCAGCCAGGCTCCTA** |
|  | pEG1.103 3R-sequence exon 10 | ACACTCTTTCCCTACACGACGCTCTTCCGATCTNNNN**TTGCGTGGAGTGTGTGGACCT** |
|  | pEG1.103 4F-sequence exon 11 | TGGAGTTCAGACGTGTGCTCTTCCGATCT**CCCATCCTGGGACAAGGCGG** |
|  | pEG1.103 4R-sequence exon 11 | ACACTCTTTCCCTACACGACGCTCTTCCGATCTNNNN**GGTTGTGGGGCCTGAGCTGT** |
|  | pNL1.243 F4- sequence exon 11 | TGGAGTTCAGACGTGTGCTCTTCCGATCT**GTGGGAGCTTGGGACACC** |
|  | pNL1.243 R4- sequence exon 11 | ACACTCTTTCCCTACACGACGCTCTTCCGATCTNNNN**CAGCACCTCTTGGGCCTG** |
|  | pNL1.243 F6- sequence exon 15 | TGGAGTTCAGACGTGTGCTCTTCCGATCT**AGTGTGTGGCTCCTGAGAGA** |
|  | pNL1.243 R6.2- sequence exon 15 | ACACTCTTTCCCTACACGACGCTCTTCCGATCTNNNN**CTATGGGTCATCCCACCTGC** |
|  | pEG1.136 1F- sequence around sgIntron-1 | ACACTCTTTCCCTACACGACGCTCTTCCGATCTNNNN**ATCAAAGAGAGACAGCACCCG** |
|  | pEG1.136 1R- sequence around sgIntron-1 | TGGAGTTCAGACGTGTGCTCTTCCGATCT**GGGCACAAGGGTACCTACG** |

**Table S3: Melting temperatures of recombinant DNMT3A2 variants by DSF**

| **Variant** | **Replicate 1 (°C)** | **Replicate 2 (°C)** |
| --- | --- | --- |
| DNMT3A2_WT | 43.4 ± 0.1 | 43.2 ± 0.1 |
| DNMT3A2_C541Y | 44.5 ± 0.2 | 44.1 ± 0.0 |
| DNMT3A2_D781N | 43.9 ± 0.1 | 43.9 ± 0.0 |
| DNMT3A2_4aaDel | 45.8 ± 1.0 | 43.2 ± 0.1 |
| DNMT3A2_G532N | 45.1 ± 0.2 | 44.7 ± 0.2 |
